# Object features survive motion at saccadic speed but lose their link to specific objects

**DOI:** 10.64898/2026.11.09.765345

**Authors:** Melis Ince, Carolin Hübner, Martin Rolfs

## Abstract

Saccadic eye movements sweep objects across the retina at high speed, and this motion is routinely omitted from conscious perception. Whether object identity survives such motion is unclear. In two preregistered experiments (N = 20 and N = 19), we asked how object motion at saccadic speeds, while the eyes fixate, affects object continuity in the object-reviewing paradigm (Kahneman et al., 1992; Sasi et al., 2023). Two Gabor patches each carried a digit, then disappeared and reappeared in place, reappeared displaced, or travelled continuously to their new locations; the intervening interval lasted 300 ms or was matched to the mean or peak speed of a saccade. A target digit then appeared in one patch, and participants identified it. At the long duration, we replicated the Object-Specific Preview Benefit (OSPB): responses were faster when the target had been previewed in that same object than in the other one. At saccadic durations, the OSPB disappeared, and adding a distinguishing surface feature in Experiment 2 did not restore it. The general preview benefit, faster responses to any previewed digit than to a new one, persisted at every duration. Object features therefore survive motion at saccadic speed, but their link to a particular object does not.

## Introduction

We explore our visual environment through saccades, rapid eye movements that redirect the fovea several times each second. Because high-acuity vision is limited to the fovea, these movements are essential for sampling a scene in detail. Each saccade, however, also displaces the retinal image, updating the retinal location of every object up to four times per second. Despite this frequent displacement, we perceive a stable and continuous world. Which information the visual system carries from one retinal location to the next, and how it keeps that information tied to the right object, is still not fully understood. Here we ask whether the information intervening such displacements— visual motion at saccadic speed—can convey object-specific information.

During a saccade, the rapid movement of the eye sweeps a smeared image across the retina, which is rarely available to conscious perception (Campbell & Wurtz, 1978). If visual information during saccades is not consciously available, is it also uninformative to perception or can it convey information? Several studies suggest that not all information contained in saccadic smear is lost. Indeed, observers can perceive motion during the saccade itself, provided the stimulus is optimized for the speeds involved (Castet, 2009; Castet et al., 2002; Castet & Masson, 2000, 2000; Garćia-Pérez & Peli, 2001; Nicolas et al., 2026). Moreover, motion smear is not erased entirely but effectively masked, largely by the post-saccadic image (Balsdon et al., 2018; Schweitzer, Doering, et al., 2025). It is also put to use: intra-saccadic motion streaks help link object locations across the saccade (Schweitzer & Rolfs, 2020), leave early neural signatures of that linking process (Schweitzer, Seel, et al., 2025), and jump-start corrective gaze behavior within tens of milliseconds of landing (Schweitzer & Rolfs, 2021). These findings suggest that saccadic smear can inform perception and behavior even when it is not available to conscious report (Higgins & Rayner, 2015; Rolfs & Schweitzer, 2022).

Similarly to saccadic smear, external motion of an object across the retina at fixation can escape conscious perception (Mattler & Fendrich, 2007, 2010; Stein et al., 2019, 2020), in particular when it follows the kinematics of saccades (Rolfs et al., 2025). Because this motion could nevertheless provide a cue of object continuity, it offers an alternative way to study how objects are tracked across saccades. In a recent study, we found that the visual system uses visual motion at saccadic speeds to track objects, even when the motion cannot be consciously detected (Ince et al., 2025). Thus, while these findings suggest that object correspondence is preserved throughout saccadic motion, they leave open whether an object’s features are carried along with it.

Kahneman et al. (1992) proposed that the visual system opens a temporary object file for each object it individuates, indexed by the object’s changing location and holding the features currently attributed to it. Because the file persists while the object moves, a feature encountered at one moment remains available at the next. The object-reviewing paradigm (Figure 1A) makes this distinction measurable (Kahneman et al., 1992). It consists of two objects placed on opposite sides of a horizontal or vertical axis before moving to new locations. In addition to their locations, the objects differ in shape and color. In the preview display, both objects contain a non-identical digit shown in their center. This is followed by a linking display, in which these digits briefly disappear as the objects move to their final locations. During the subsequent target display, once the objects reach their final locations, one digit (target) appears in only one of these objects. The target is either the digit that this object itself contained during the preview (same-object condition), the digit that the other object contained (different-object condition), or a digit that was contained in neither of them. The latter condition (no match condition) provides a baseline in which no preview information is available. Participants’ task is to verbally report the presented digit in the target display as fast and accurately as possible. If the object-specific information has travelled with it, the digit it carried should still be bound to it, and naming should be faster in the same-object than in the different-object condition. This difference, RT_different_ minus RT_same_, is the ‘object-specific preview benefit’ (OSPB). The no-match condition supplies a second, more general measure of preview retention: the advantage of a previewed digit over one that appeared in neither object, RT_no-match_ minus RT_different_, which we call the general preview benefit (GPB). Thus, while the GPB requires only that a digit was seen, the OSPB requires that it still be bound to a particular object. Henderson & Anes (1994) drew the same distinction between object-file review (here, OSPB) and type priming (here, GPB) in visual identification across eye fixations.

**Figure 1.**
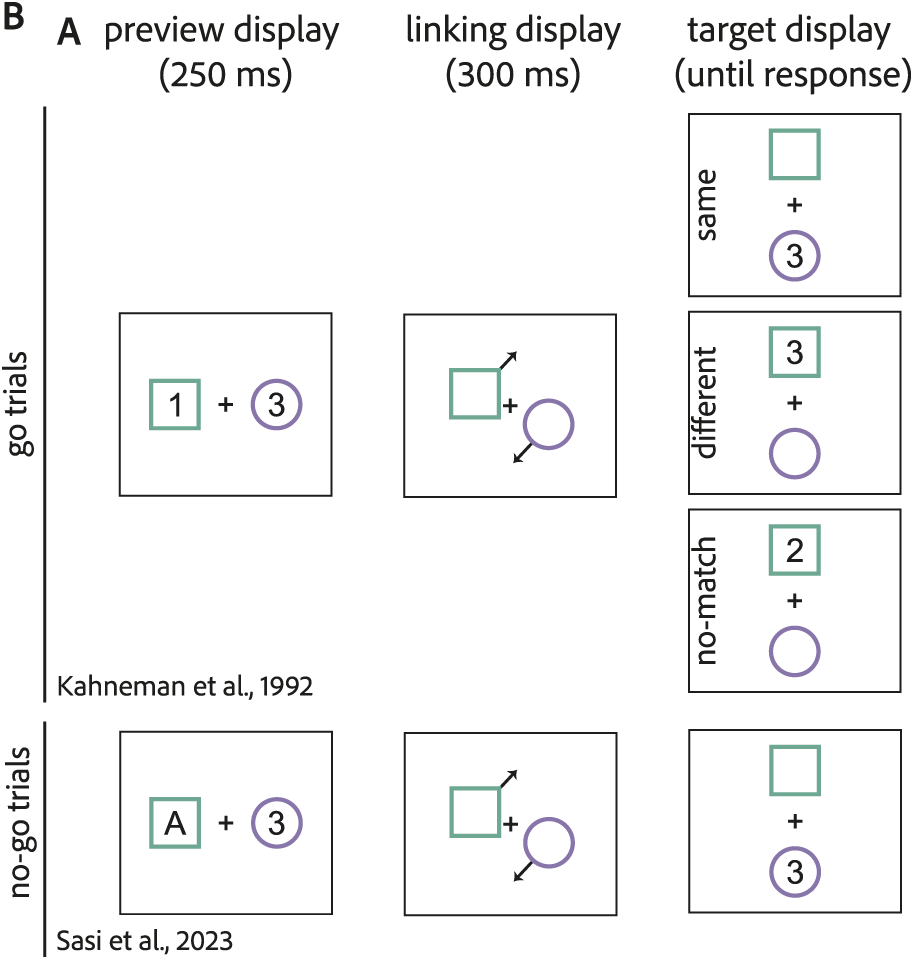
The original (Kahneman et al., 1992) and the modified (Sasi et al., 2023) object-reviewing paradigms. **A.** Two objects with distinct visual properties (shape and color) are shown in the preview display with non-identical digits presented in their centers. In the linking display, the digits disappear and the objects move to their new locations. Finally, on the target display, only one digit appears in one of the objects. The identity of this digit and its location differ depending on the object condition in the trial (same, different, or no-match). Participants are asked to report the target digit as fast as possible. **B.** In a recently modified version, Sasi et al. (2023) added ‘go/no-go trials’ to the paradigm. While ‘go trials’ are the same object conditions as in the original paradigm, ‘no-go trials’ start with a letter and a digit on the preview display, and participants are required to press the space bar instead of reporting the target digit at the end of the trial.

The original object files paradigm, however, featured a potential caveat: the task did not require participants to pay attention to the preview display. This may have contributed to object-specific preview benefits that were often small and reportedly not robust (Noles et al., 2005). Noles et al., (2005) addressed this by introducing the modified reviewing paradigm, requiring an explicit match/no-match judgment on the target letter with respect to preview letters. The modified paradigm found larger and more reliable OSPBs than the original version. But the modified task also turned the paradigm into a visual working memory task rather than a purely perceptual one.

Recently, Sasi et al. (2023) removed this issue by adding catch trials to the original paradigm, which would encourage participants to attend to the preview display without introducing a memory component. In their version, “go trials” were similar to the original paradigm. Two digits were presented in the preview display, and the participants reported the target via keypress. Critically, the design contained “no-go trials”, in which the preview display included a letter and a digit, and participants were asked to refrain from reporting the target and instead press the space bar at the end of the trial (Figure 1B). This carefully modified go/no-go reviewing paradigm successfully replicated the OSPB, providing a new, reliable way to test object files in visual perception. In our study, we followed the same approach to ensure participants’ attention to the preview display.

### The present study

In two pre-registered experiments (Experiment 1: https://osf.io/3zj6v; Experiment 2: https://osf.io/wjp5s), we asked whether object-specific information is conveyed by visual motion at saccadic speeds—that is, when an object is moving as fast as a saccade would move the retinal image. To address this question, we adapted the go/no-go reviewing paradigm of Sasi et al. (2023) in three ways.

First, and unlike most object-file paradigms, we used two identical Gabor patches as objects in Experiment 1 (Figure 2). Gabors can be displaced continuously at high speed without producing a visible motion streak (Rolfs et al., 2025; for stimulus details, see Methods). Because the two objects are physically identical, the only information that distinguishes them is their spatiotemporal history and the digit each contained during the preview. In Experiment 2, we added a light and a dark grey ring around the two objects during the preview and target displays, giving the objects a distinguishing surface feature. This manipulation addressed whether surface information can support object-specific benefits in comparison to spatiotemporal information alone.

**Figure 2.**
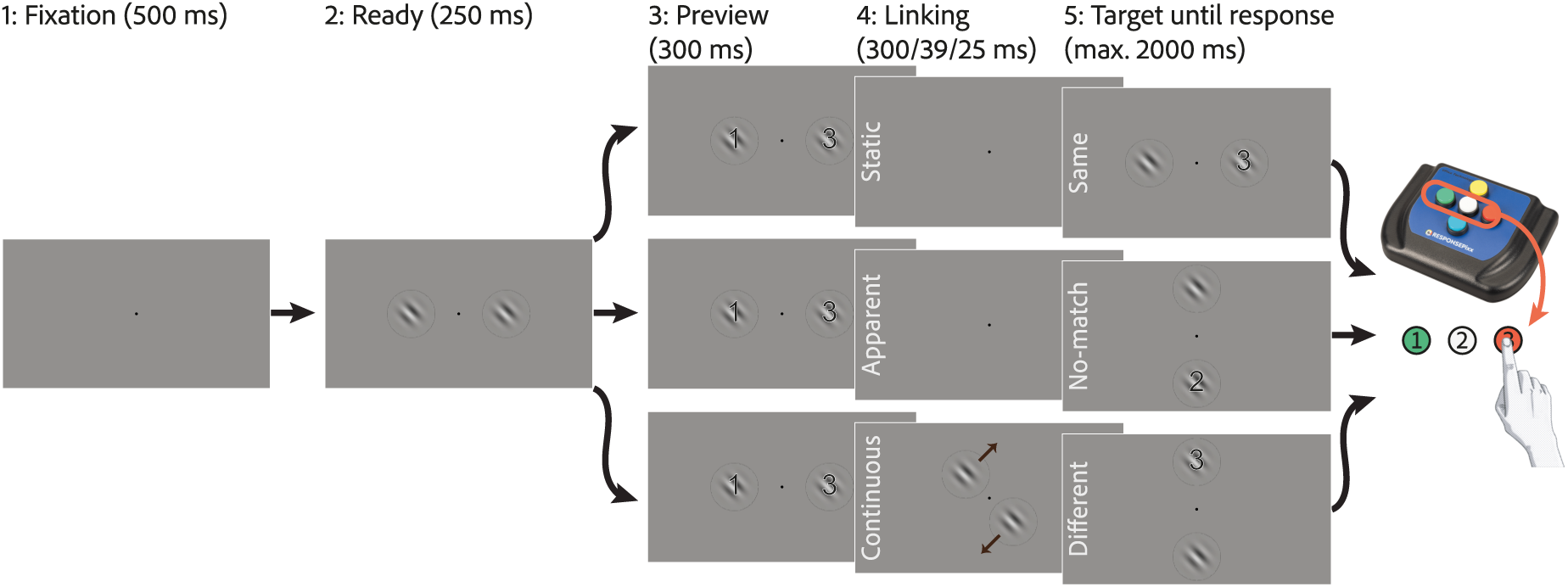
Trial sequence in Experiment 1. Each trial began with a fixation display, during which participants were asked to fixate at the center of the screen throughout the trial. In the “ready” display, two Gabor patches appeared either horizontally or vertically aligned. In the preview, participants either saw two different digits (go trials) or a digit and a letter (no-go trials) within the Gabor patches. In the linking display, depending on the trial’s motion condition, there could be a blank display (static and apparent) or continuous motion (continuous). The duration of the linking display depended on the duration condition and could be 300 (D_long_), ∼39 (D_vsac_), or 25 (D_vpeak_) ms long. In the target display, one digit either appeared within the same object as it was in the preview (as shown in the first row, same-object) or in the other object (as shown in the third row, different-object). Alternatively, a new digit could appear in either object (as shown in the second row, no-match). Experiment 2 used the same sequence, with a light or dark grey ring added around each Gabor patch in the preview and target displays.

Second, we implemented three motion conditions within the linking display that differed in the spatiotemporal cues available to link the preview and the target displays. In the static condition, the objects disappeared and reappeared in the same locations; nothing was displaced, and object identity remained fully specified by location. In the apparent condition, objects disappeared and reappeared in new locations with no motion information in between; because each preview location was equidistant from both target locations, the correspondence between preview and target objects was ambiguous (no-consistent apparent motion). In the continuous condition, the objects travelled along their trajectories, so that object correspondence was specified unambiguously by visual motion. The static and apparent conditions thus bracket the continuous one: the first offers correspondence without displacement, the second offers displacement without consistent correspondence.

Third, we manipulated the duration of the linking display while holding motion distance constant, thereby determining motion speed. The manipulation also applies to displays in which nothing moves: in the continuous condition, the linking display duration sets the objects’ speed, whereas elsewhere it sets the duration of the blank interval. Experiment 1 used three durations: a long duration (*D_lon_*_g_, 300 ms) replicating Sasi et al. (2023), which anchors our design to the established effect, and two short durations corresponding to the time required to traverse 6 degrees of visual angle (dva) at the mean velocity (*D_vsac_*, ∼39 ms) and the peak velocity (*D_vpeak_*, ∼25 ms) of a saccade of that amplitude.

For the long duration, we expected to replicate the standard effect: an OSPB in the static and continuous conditions, and none in the apparent condition, where correspondence is ambiguous. The critical test is the continuous condition at saccadic durations, for which we consider two competing hypotheses: If object-specific preview information survives motion at saccadic speeds, the OSPB should persist there. If only object correspondence survives, but not the object’s features, the OSPB should vanish while the GPB remains. Importantly, our previous finding that objects can be tracked through such motion only establishes the former (Ince et al., 2025). Finding only a GPB would indicate that preview information is retained but no longer assigned to the object that carried it. Experiment 2 adds a third possibility: that a distinguishing surface feature enables the object-specific benefit where spatiotemporal information alone may not.

Independently of the two preview benefits, we predicted that continuous-motion conditions would yield shorter response times than apparent-motion conditions overall. This is because continuous motion allows the visual system to maintain object correspondence across locations while apparent motion requires correspondence to be re-established at the final location, adding an additional processing step.

## Experiment 1

### Methods

#### Participants

We recruited twenty healthy human participants (14 females and 6 males, 18 right-handed, 12 right-eye dominant), aged 18-35 (Median = 26), through word of mouth (including members of the Active Perception and Cognition group) and the local participant pool (SONA) at the Department of Psychology of Humboldt-Universität zu Berlin, which is open to both student and non-student populations. All participants were naïve to the study. They provided informed consent before testing and the project was approved by the Department’s Ethics Committee (reference number 2019-08). Participants were compensated via accreditation of work hours, payment of 10€/hour, or course credit, at their choosing. As pre-registered, data collection continued until complete datasets were obtained from 20 participants; participants who did not complete all experimental sessions were excluded from analyses and replaced.

Before data collection, we assessed ocular dominance using the Miles test (Miles, 1930) to determine which eye would be tracked during the experiment and tested participants’ visual acuity using a Snellen chart embedded in a Polatest vision testing instrument (Zeiss, Oberkochen, Germany). All participants had normal (n=13) or corrected-to-normal (n=7) visual acuity.

#### Equipment

Stimuli were projected onto a 220 x 124 cm screen (Celexon HomeCinema High Contrast Frame, 100” – Dynamic Slate ALR, Tharston, Norwich, UK) using a gamma-linearized PROPixx DLP Projector (Vpixx Technologies, Saint-Bruno, QC, Canada) with a spatial resolution of 960 x 540 pixels (greyscale) and a refresh rate of 1440 Hz. The experimental code was implemented in MATLAB (MathWorks, Natick, MA, USA), using the Psychophysics toolbox (Kleiner et al., 2007), running on a Dell XPS 8940 operating on Ubuntu 20.04.6 LTS. Eye movements were recorded with an EyeLink 1000+ system (SR Research, Ottawa, ON, Canada) at a sampling rate of 1000 Hz, with the head stabilized by a chin and forehead rest at a viewing distance of 270 cm. Responses were collected using a ResponsePixx HANDHELD (VPixx Technologies Inc., Saint-Bruno, QC, Canada), a five-button response device with buttons arranged in a plus sign (see Figure 2). Only the three horizontally aligned buttons were used; the top and bottom buttons were disabled. Restricting responses to a single row equated the movement required for each of the three digits, so that response times cannot differ simply because one button is easier to reach than another. During the experiment, the three active buttons were illuminated to indicate the available responses and avoid confusion. Possible target digits (1, 2, 3) were assigned from the left to the right button in ascending order.

#### Stimuli

On each trial, two Gabor patches (spatial frequency of 1 cycle per dva [cyc/dva]; full contrast; sigma: 1/3 dva) were displayed. In the continuous motion trials, the Gabor patches moved along a diagonal path, perpendicular to the Gabor’s orientation (i.e., along the axis of luminance modulation). The Gabor patches were placed at a horizontal distance of 4.24 dva from the fixation point on the preview display and at a vertical distance of 4.24 dva from the fixation point on the target display, or vice versa (see example for horizontal to vertical in Figure 2). Their resulting motion trajectory spanned 6 dva. Each preview location was therefore equidistant (6 dva) from both target locations, so that the geometry alone never favored one preview-to-target mapping over the other.

The digits ({1, 2, 3}) and letters ({A, C, D, H, K, M, W}) were previously rendered in Adobe Illustrator (Adobe Inc., San Jose, CA) in black Arial font with a fine white outline, exported as PNG files, and imported into MATLAB for display within a 1.9 dva x 1.9 dva bounding box. The purpose of the fine white outline was to increase readability of the letters/digits over Gabor patches.

#### Design and Procedure

We applied a within-subject, repeated-measures design. Each participant completed three experimental sessions (approximately 90 minutes each); only the first session was preceded by a brief training session (approximately 10 minutes). During training, only two linking display durations were presented to familiarize participants with the task: the long duration (300 ms) and the duration matching the saccadic mean speed (∼39 ms).

Each experimental session consisted of 9 blocks of 120 trials, resulting in a total of 3,240 trials per participant over the three sessions. The design crossed three factors: the duration of the linking display, the motion condition, and the object condition (Figure 2). Motion conditions were blocked, with block order counterbalanced across participants. Linking display durations and object conditions were interleaved within each block: every block contained the same number of trials of each combination, and their order within the block was randomized. Starting position (horizontal vs vertical object arrangement in the preview display), motion direction (clockwise vs counterclockwise), the assignment of digits to objects during the preview display, and object conditions were balanced across trials, but appeared in a random sequence.

We tracked the dominant eye throughout the trial and controlled gaze position online. Trials in which fixation was not maintained within a 2 dva radius around the fixation point, and trials in which no response was registered within 2000 ms of target appearance, were cancelled online and repeated at the end of the block. In these cases, we displayed “Fixate please” and “Please respond faster” messages, respectively. On average, 10.9% (SD = 5.1%) of trials were repeated for these reasons.

Each trial started with a blank display showing only a fixation point at the center for 500 ms. Subsequently, a “ready” display presented two Gabor patches aligned horizontally or vertically for 250 ms. Next, the preview display showed a digit or letter at the center of each Gabor patch for 300 ms. The following linking display on each trial was assigned one of three duration conditions: a long duration (*D_lon_*_g_: 300 ms, 20 deg/s, in continuous motion) and two durations matching the saccadic mean and peak velocities (*D_vsac_*: ∼39 ms and *D_vpeak_*: ∼25 ms, respectively). *D_lon_*_g_ was chosen to match the linking display duration used by Sasi et al. (2023). The two shorter durations were calculated to match the mean (*v_sac_*; 154.3 deg/s) and peak (*v_peak_*; 239.4 deg/s) saccadic speeds for the distance between the initial and final positions (6 dva). Following previous studies (Ince et al., 2025; Rolfs et al., 2025), we calculated these durations according to the main sequence derived from Collewijn et al. (1988) and Van Opstal & Van Gisbergen (1987). The fixation point remained visible throughout the trial and no additional fixed inter-trial interval was used (ITI = 0 ms).

The experiment had three motion conditions that changed how the Gabor patches behaved between preview and target displays. In the static condition, the preview and the target positions were kept the same and Gabors were blanked during the linking display. Similarly, in the apparent condition, Gabors were blanked during the linking display but they reappeared in their new locations in the target display, creating the perception of apparent motion. The continuous condition required the Gabor patches to move continuously from their initial to final locations. The linking display meant something different in each motion condition: in the continuous condition it set the objects’ speed, whereas in the static and apparent conditions it set the duration of the blank, leaving all three motion conditions matched in total trial duration.

In go trials, two digits were shown on the Gabor patches for the duration of the preview display (300 ms), and the target display then showed a single digit on one of them. In the same-object condition, the target digit matched the digit previewed on that same object, whereas in the different-object condition, it matched the digit previewed on the other object. The displayed digits were each randomly selected without replacement from their respective pools so that the two digits displayed in the preview were never identical. In the no-match condition, a new digit (the remaining third digit in the pool) appeared on one of the two objects, creating no correspondence between the preview and target stimuli. Same– and different-object conditions had equal likelihood, occurring in 20% of trials each, and the no-match condition occurred in 40% of trials. The target display remained on screen until a response was made, or for a maximum of 2000 ms. Participants reported the observed digit by pressing the corresponding key on a ResponsePixx button box (see Equipment).

The remaining 20% of trials (catch-trials) were called no-go trials, which served as an attentional control. In these trials, one letter and one digit were shown during the preview display and participants were instructed to press the space bar at the target display (which was identical to that in go trials) and refrain from reporting any digit. Participants correctly withheld a digit response on 97.0% (SD = 3.4%) of the no-go trials, confirming that they attended to the preview display throughout the experiment.

#### Data Analysis

We calculated the proportion of repeated trials by analyzing the EDF event logs. No additional offline eye-movement analysis was performed, as the online monitoring and repetition of trials made this unnecessary.

We measured response time (RT) in milliseconds as the primary dependent variable, defined as the time difference between stimulus offset and keypress. Error rate was measured as the percentage of incorrect responses across trials, providing an additional measure of task performance and attention to the experimental stimuli.

All statistical analyses were performed using MATLAB (R2024b). RT data were pre-processed following Sasi et al. (2023), with one deviation noted below. First, error trials (incorrect responses in go trials) and trials with anticipatory RTs less than 200 ms were excluded. Unlike Sasi et al. (2023), we did not apply an upper absolute RT cut-off to avoid over-cleaning the data; slow responses were instead handled by the per-condition trimming below. Within each participant and experimental condition (linking display duration condition × motion condition × object condition), trials differing from the condition mean by more than 2 SDs were removed. After excluding anticipatory responses (< 200 ms), 60,215 of the 64,800 trials (92.9%) were retained; condition-specific outlier removal (±2 SD from the condition mean) then removed a further 2,150 trials, leaving 58,065 trials (89.6%). Following outlier removal, all RTs were log-transformed for statistical analysis. Response times, mean differences, and figure values are reported as arithmetic means in milliseconds for ease of interpretation, whereas all inferential statistics (t-tests, Bayes factors, and effect sizes) were computed on log-transformed RTs. Effect sizes for planned binary comparisons are reported as Cohen’s d_z_ for paired samples, computed on the log scale.

Error rates were calculated separately for go and no-go trials. For the no-go trials, the error rate was the overall percentage of trials on which participants failed to press the space bar, serving as a measure of attentiveness to the preview display. For the go trials, error rates were calculated within each experimental condition (linking display duration condition × motion condition × object condition).

Log-transformed RTs were submitted to a 3 × 3 × 3 repeated-measures ANOVA with linking display duration condition, motion condition, and object condition as within-participant factors. Because each factor had three levels and the assumption of sphericity was violated in some conditions, Greenhouse-Geisser corrections were applied to the degrees of freedom, and we report Greenhouse-Geisser corrected degrees of freedom and p-values together with partial eta-squared (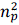) as a measure of effect size.

Following the rmANOVA, we used planned comparisons to test the preview benefits. As defined in the Introduction, the object-specific preview benefit (OSPB) was calculated as the difference in mean RT between different-object and same-object trials, and the general preview benefit (GPB) as the difference between no-match and different-object trials. Comparisons were conducted using paired-samples t-tests. Because we had a directional prediction for the OSPB (faster responses on same-than different-object trials), OSPB t-tests were one-tailed. General preview benefits, for which we had no directional prediction, were tested two-tailed. The p-values from planned comparisons were corrected for multiple comparisons using the Benjamini-Hochberg false discovery rate procedure (Benjamini & Hochberg, 1995), implemented via the mafdr function (MATLAB Bioinformatics Toolbox). FDR correction was applied separately within each family of comparisons rather than across all tests, so that each correction controlled the false discovery rate for one hypothesis.

An equivalent 3 × 3 × 3 repeated-measures ANOVA, with the same factors and Greenhouse-Geisser correction, was conducted on go-trial error rates to assess whether the RT effects were mirrored in accuracy and to rule out speed-accuracy trade-offs.

For each planned comparison we additionally computed Bayes factors using the bayesFactor toolbox (Krekelberg, 2024) with a default Cauchy prior (scale r = 0.707) on the standardized effect size. The directionality of each Bayes factor matched the corresponding t-test: one-tailed for OSPB comparisons and two-tailed for GPB comparisons. We write BF_+0_ for comparisons tested one-tailed against the predicted direction (the OSPBs) and BF_10_ for those tested two-tailed (the GPBs, and the exploratory tests reported in the Results).

## Results

Log-transformed RTs were analyzed using a 3 × 3 × 3 repeated-measures ANOVA with linking display duration, motion condition, and object condition as within-participant factors. Mauchly’s test indicated that the assumption of sphericity was violated for the main effects of linking display duration (χ²(2) = 18.72, p < .001) and object condition (χ²(2) = 21.74, p < .001), and for the linking display duration × motion condition × object condition interaction (χ²(35) = 55.86, p = .014). Greenhouse-Geisser corrected degrees of freedom and p values are therefore reported for all effects.

There were significant main effects of linking display duration, F(1.21, 23.08) = 149.37, p < .001, 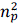 = .89, motion condition, F(1.90, 36.13) = 12.10, p < .001, 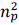 = .39, and object condition, F(1.18, 22.34) = 5.49, p = .024, 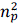 = .22. These were qualified by significant interactions of linking display duration with motion, F(3.48, 66.08) = 19.51, p < .001, 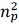 = .51, and linking display duration with object, F(2.93, 55.61) = 15.19, p < .001, 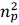 = .44, a motion-by-object interaction, F(3.07, 58.30) = 3.31, p = .025, 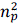 = .15, and a three-way linking display duration × motion condition × object condition interaction, F(4.70, 89.26) = 4.29, p = .002, 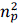 = .18. The effect of object condition and its interactions with linking display duration and motion indicate that preview effects varied across conditions, motivating the planned comparisons reported below, which separate these findings into object-specific and general preview benefits at each linking display duration and motion condition.

The same 3 × 3 × 3 repeated-measures ANOVA on go-trial error rates for a potential speed-accuracy trade-off showed no significant difference. There were no main effects of linking display duration, F(1.12, 21.31) = 3.15, p = .086, 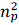 = .14, motion condition, F(1.77, 33.72) = 0.33, p = .693, 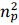 = .02, or object condition, F(1.04, 19.67) = 0.15, p = .711, 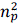 = .01, and none of the interactions reached significance (all p > .08; linking display duration × object, F(2.64, 50.10) = 2.46, p = .081, 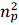 = .12).

### Object-Specific Preview Benefit (RT_different_ – RT_same_)

We conducted planned comparisons to test for OSPBs across all linking display duration and motion conditions (see Figure 3A). As expected, at *D_lon_*_g_, we found significant OSPBs in both the static condition (Mean±95%CI: 8.5±7.2 ms, t(19) = 3.12, p_fdr_ = .013, Cohen’s d_z_ = 0.697, BF_+0_ = 16.50) and the continuous condition (18.3±8.6 ms, t(19) = 5.09, p_fdr_ < .001, d_z_ = 1.138, BF_+0_ = 818.58), while the apparent condition showed no significant OSPB (5.1±7.6 ms, t(19) = 1.00, p_fdr_ = .497, d_z_ = 0.223, BF_+0_ = 0.60).

**Figure 3.**
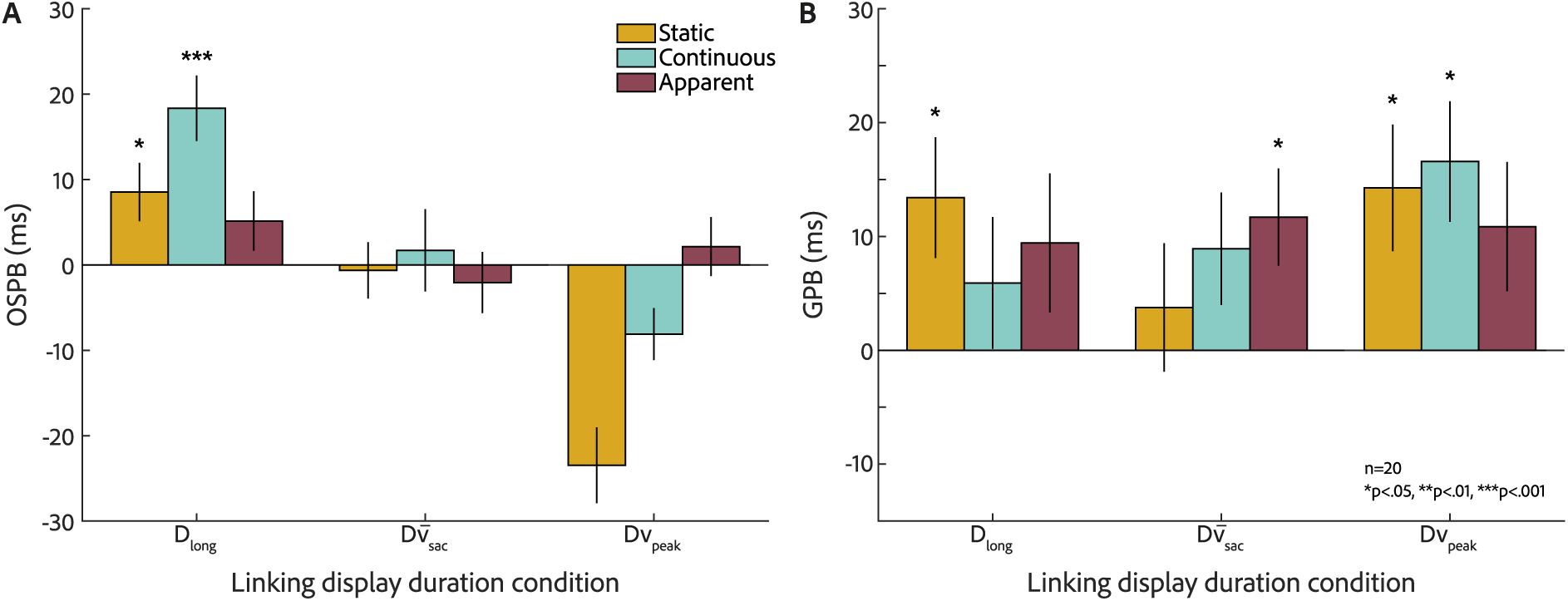
Experiment 1: Object-specific (OSPB) and general preview benefits (GPBs) shown for each motion and linking display duration condition. Asterisks mark comparisons that remained significant after Benjamini-Hochberg FDR correction, with thresholds shown on the lower bottom; OSPB tests were one-tailed and GPB tests two-tailed. The colors indicate motion conditions, and the x-axis shows linking display durations. Error bars represent bootstrapped standard errors of the mean (1,000 resamples). **A.** OSPBs calculated as RT_different_ – RT_same_, and their arithmetic mean is shown in milliseconds. **B.** General preview benefits (GPBs) calculated as RT_nomatch_–RT_different_ and their arithmetic mean is shown in milliseconds.

Contrary to our main hypotheses, at *D_vsac_*, no significant OSPBs were observed across any motion condition (static: –0.6±7.2 ms, t(19) = 0.20, p_fdr_ = .632, BF_+0_ = 0.27; continuous: 1.7±10.6 ms, t(19) = 0.33, p_fdr_ = .632, BF_+0_ = 0.31; apparent: –2.1±7.7 ms, t(19) = –0.60, p_fdr_ = .929, BF_+0_ = 0.15).

At *D_vpeak_*, responses were faster on different-than on same-object trials in the static (– 23.5±9.4 ms, t(19) = –4.95, d_z_ = –1.106) and continuous (-8.1±6.5 ms, t(19) = –2.33, d_z_ = – 0.521) conditions. Our planned comparisons were one-tailed in the predicted direction and cannot evaluate differences of this sign (both p_fdr_ = 1.00; BF_+0_ = 0.03 and 0.06, respectively). Treated as exploratory and tested two-tailed, the static difference is substantial (p < .001, BF_10_ = 306.26), whereas the continuous difference is weak (p = .031, BF_10_ = 2.03; both uncorrected). The apparent condition showed no effect (2.1±7.6 ms, t(19) = 0.49, p_fdr_ = .632, d_z_ = 0.111, BF_+0_ = 0.36).

Error rates did not follow the OSPB pattern of significant differences between conditions. At *D_lon_*_g_, errors were 5.9% (SD = 3.8%) for same-, 7.1% (SD = 5.4%) for different-, and 7.2% (SD = 9.0%) for no-match trials; at *D_vsac_*, 5.8% (SD = 3.9%), 5.4% (SD = 4.5%), and 6.1% (SD = 8.8%); and at *D_vpeak_*, 6.1% (SD = 4.3%), 5.3% (SD = 3.5%), and 6.1% (SD = 9.1%).

### General Preview Benefit (RT_no-match_ – RT_different_)

At *D_lon_*_g_, we found a GPB in the static condition (13.4±11.3 ms, t(19) = 2.60, p_fdr_ = .043, d_z_ = 0.581, BF_10_ = 3.19), while continuous (5.9±12.4 ms, t(19) = 1.14, p_fdr_ = .301, BF_10_ = 0.41) and apparent (9.4±13.8 ms, t(19) = 1.61, p_fdr_ = .160, BF_10_ = 0.70) conditions showed none. At *D_vsac_*, the apparent condition showed a significant GPB (11.7±9.2 ms, t(19) = 2.98, p_fdr_ = .035, d_z_ = 0.665, BF_10_ = 6.33), while the continuous (8.9±10.7 ms, t(19) = 2.06, p_fdr_ = .096, BF_10_ = 1.32), and the static (3.8±12.4 ms, t(19) = 0.64, p_fdr_ = .529, BF_10_ = 0.28) were not significant. At *D_vpeak_*, the static (14.3±11.8 ms, t(19) = 2.56, p_fdr_ = .043, d_z_ = 0.573, BF_10_ = 3.00) and continuous (16.6±11.7 ms, t(19) = 3.18, p_fdr_ = .035, d_z_ = 0.711, BF_10_ = 9.31) conditions showed significant GPBs, while the apparent condition did not (10.9±12.3 ms, t(19) = 1.89, p_fdr_ = .112, BF_10_ = 1.02). These findings suggest that GPBs, independent of object identity, persist across linking display durations, particularly at shorter durations where object-specific benefits are absent (Figure 3B).

Preview benefits are difference scores computed within a motion condition and thus do not speak to the cost of establishing correspondence in the first place. Continuous motion never interrupts spatiotemporal continuity, whereas apparent motion requires correspondence to be re-established at the target location; if that re-establishment carries a processing cost, overall response times should be faster in the continuous than in the apparent condition. We therefore compared the two conditions directly at each duration. We ran two-tailed paired t-tests for each linking display duration. At *D_lon_*_g_, RTs in the continuous condition were significantly slower than in the apparent condition (12.7 ms difference, t(19) = 4.29, p_raw_ < .001, d_z_ = 0.960), indicating a significant effect in the opposite direction of our hypothesis. At the two saccadic durations, we found no significant differences: *D_vsac_* showed a 6.1 ms difference (t(19) = 1.85, p_raw_ = .079, d_z_ = 0.414), and *D_vpeak_* showed a 5.1 ms difference (t(19) = 1.18, p_raw_ = .252, d_z_ = 0.264).

## Discussion

We found OSPBs for the long linking display duration in both the static and continuous conditions, confirming their robustness when sufficient processing time is available (consistent with Sasi et al., 2023). Because the objects do not change location in the static condition, the object-specific claim rests on the continuous condition, where object and location dissociate. Obtaining that benefit with identical Gabor patches indicates that objects do not need to differ in shape or color for object-specific feature transfer. As expected from a perceptually ambiguous condition, we found no OSPB in our apparent motion condition.

Neither saccadic duration produced an OSPB. At *D_vsac_*, no significant OSPBs emerged in any motion condition; at *D_vpeak_*, responses were instead faster on different-than on same-object trials in the static and, less markedly, the continuous condition. Because our planned comparisons were one-tailed in the predicted direction, they cannot evaluate a difference of this sign, and we treat the reversal as exploratory. While it might be due to our experimental design (see General Discussion), these findings suggest limitations in the visual system’s ability to maintain object-specific information during saccade-like motion.

Furthermore, we found GPBs, particularly at saccadic durations, indicating that the visual system still has access to the target previews regardless, but can no longer connect them to specific objects or locations.

We had predicted faster responses in the continuous than in the apparent condition, since continuous motion maintains object correspondence whereas apparent motion requires it to be re-established. We found the opposite: responses were slower in the continuous condition, significantly so at the long linking duration. The benefit of uninterrupted correspondence may be offset by the cost of tracking a moving object through the linking display, a process that is resource-limited (Holcombe & Chen, 2012). Such a cost should grow with the duration of the motion, and the difference between the two conditions did decrease across the three durations. This account is post hoc, and it leaves the preview benefits untouched, since those are computed within each motion condition. Moreover, as reported below, the slowing did not reach significance in Experiment 2.

While we calculated *D_vpeak_* based on saccade kinematics (Collewijn et al., 1988; Van Opstal & Van Gisbergen, 1987), but our objects moved at constant velocity rather than accelerating to a peak and decelerating again as the eyes do. A real saccade therefore spends only a moment of its trajectory at peak velocity (*v_sac_*), and its mean velocity is correspondingly lower. A duration matched to peak velocity has no clear counterpart in an actual eye movement, so Experiment 2, which again used constant-velocity motion, retained only *D_lon_*_g_ and *D_vsac_* (representing the average saccade duration) for the linking display duration.

Finally, spatiotemporal continuity alone proved sufficient to produce OSPBs, without any distinguishing features between Gabor patches. In Experiment 2 we nevertheless added dark and light grey rings around the objects in the preview and target displays, to test whether a surface feature would strengthen the preview benefits or extend it to saccadic durations.

## Experiment 2

### Methods

#### Participants

We recruited twenty healthy human participants and included data from nineteen in the final analysis (17 females, 1 male, and 1 non-binary; 17 right-handed, 11 right-eye dominant), aged 19 to 32 (Median = 23). One participant was excluded prior to data analysis for falling outside the preregistered age range (18 to 35). Recruitment had already ended upon reaching the target sample size of 20 participants, and the age-eligibility violation was identified only afterward; consequently, the excluded participant was not replaced. All participants had normal (n = 6) or corrected-to-normal (n = 13) vision, with a visual acuity score of 20/20.

We ensured that all participants were naïve to the study by not recruiting previously tested participants. Participant recruitment, compensation, pre-testing procedures, informed consent, and ethical approval were identical to Experiment 1.

#### Stimuli and Equipment

Equipment was identical to Experiment 1. In the preview and target displays, we added a light and dark grey ring around each Gabor patch, giving the objects a distinguishing surface feature. The light grey ring had a normalized luminance of 0.80 (RGB: 204, 204, 204) and the dark grey ring 0.20 (RGB: 51, 51, 51), both defined against the mid-grey background (normalized luminance: 0.50), such that the two rings were opposite in contrast polarity. The placement of the light and dark grey rings was randomized across all trials, but each Gabor patch kept the same ring for the duration of a trial, so that the two objects were always distinguishable by contrast polarity. The rings followed the same temporal timing as the digits/letters in the preview and target displays, and were not shown during the continuous motion to prevent visible motion streaks. All other stimuli were identical to Experiment 1.

#### Design and Procedure

The experimental design was identical to Experiment 1, with the following modifications. Only two of the three linking display duration conditions were included: the long duration, *D_lon_*_g_, and the duration matching the mean saccadic speed, *D_vsac_*.

Each participant completed two experimental sessions (approximately 90 minutes each) instead of three. As in Experiment 1, only the first session was preceded by a brief training session (approximately 10 minutes), consisting of 5 blocks of 10 trials each. The two linking display durations used in the main experiment were presented during training. Each experimental session consisted of 12 blocks of 90 trials each, for a total of 2,160 trials per participant across the entire experiment.

All other aspects of the design, including motion conditions, counterbalancing, blocked and interleaved conditions, trial cancellation criteria, and fixation monitoring, were identical to Experiment 1. On average, 10.8% (SD = 5.8%) of trials were repeated due to fixation breaks or responses exceeding the 2000 ms response window.

#### Trial Procedure

The trial procedure was identical to Experiment 1.

#### Data Analysis

Data analysis followed the same pipeline as Experiment 1. After excluding anticipatory responses (< 200 ms), 38,101 of the 41,040 trials (92.8%) were retained; condition-specific outlier removal (±2 SD from the condition mean) then removed a further 1,356 trials, leaving 36,745 trials (89.5%) after RT-based trimming.

Because Experiment 2 included only two linking display duration conditions, log-transformed RTs were submitted to a 2 × 3 × 3 repeated-measures ANOVA with linking display duration, motion condition, and object condition as within-participant factors. Greenhouse-Geisser corrected degrees of freedom and p-values are reported for all effects. As in Experiment 1, FDR correction was applied separately within each family of comparisons, here across 6 tests (2 linking display durations × 3 motion conditions) rather than 9. All other analysis decisions, including log transformation, outlier removal, effect size measures, and Bayes factor computation were identical to Experiment 1.

## Results

Log-transformed RTs were analyzed using a 2 × 3 × 3 repeated-measures ANOVA with linking display duration, motion condition, and object condition as within-participant factors. Mauchly’s test indicated that the assumption of sphericity was violated for the main effect of object condition (χ²(2) = 6.49, p = .039, ε = .759). The linking display duration had only two levels and is therefore exempt from the sphericity assumption.

There were significant main effects of linking display duration, F(1, 18) = 216.93, p < .001, 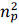 = .923, and object condition, F(1.52, 27.33) = 11.97, p < .001, 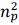 = .399, but no significant main effect of motion condition, F(2.00, 35.91) = 0.48, p = .621, 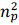 = .026. These were qualified by significant interactions of linking display duration with motion condition, F(1.97, 35.48) = 6.45, p = .004, 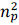 = .264, linking display duration with object condition, F(1.70, 30.61) = 19.04, p < .001, 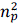 = .514, a motion-by-object interaction, F(2.94, 52.84) = 4.63, p = .006, 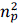 = .205, and a three-way linking display duration × motion condition × object condition interaction, F(3.15, 56.79) = 4.32, p = .007, 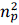 = .194. As in Experiment 1, the effect of object condition and its interactions with linking display duration and motion indicate that preview effects varied across conditions, motivating the planned comparisons reported below.

We conducted the same 2 × 3 × 3 repeated-measures ANOVA on go-trial error rates. Mauchly’s test indicated no significant violations of sphericity for any of the effects in the error-rate ANOVA (all p > .11). We observed a significant main effect of linking display duration, F(1, 18) = 5.96, p = .025, 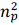 = .249, reflecting overall lower error rates at the mean saccadic duration than at the long duration, and a significant linking display duration × motion condition interaction, F(1.63, 29.36) = 5.10, p = .017, 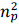 = .221. Critically, there were no significant effects of object condition or any interaction involving object condition. Planned comparisons on error rates between same– and different-object trials were non-significant in all conditions, ruling out a speed-accuracy trade-off.

### Object-Specific Preview Benefit (RT_different_ – RT_same_)

We conducted planned comparisons to test for OSPBs across all linking display duration and motion conditions (see Figure 4A). At *D_lon_*_g_, we found a significant OSPB in the continuous condition (21.1±10.6 ms, t(18) = 4.33, p_fdr_ = .001, d_z_ = 0.993, BF_+0_ = 163.37), while the static (3.5±11.1 ms, t(18) = 1.01, p_fdr_ = .491, d_z_ = 0.231, BF_+0_ = 0.620) and apparent (1.2±8.2 ms, t(18) = 0.35, p_fdr_ = .545, d_z_ = 0.081, BF_+0_ = 0.320) conditions showed no significant benefit.

**Figure 4.**
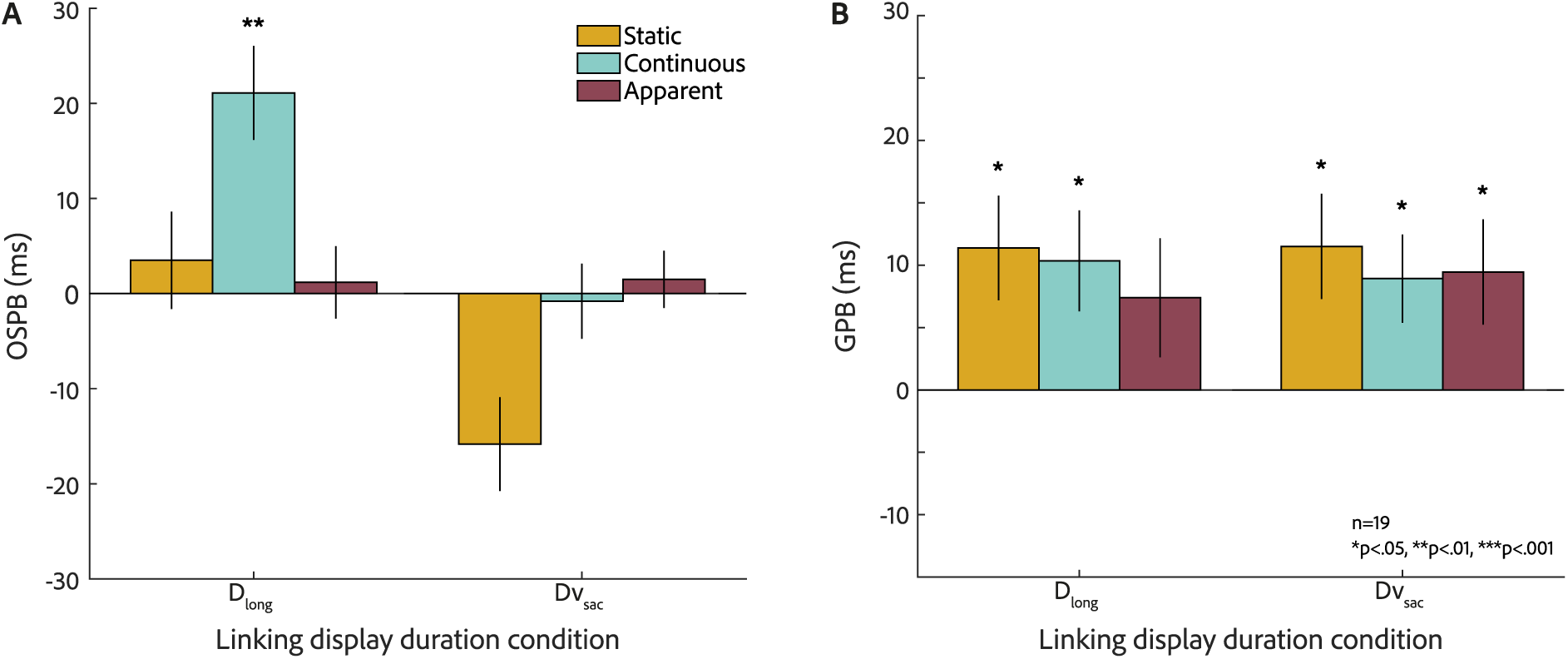
OSPB and GPB results of Experiment 2. Conventions as in Figure 3, with the exception that only two linking display durations were tested, *D_lon_*_g_ and *D_vsac_*.

At *D_vsac_*, no significant OSPBs were observed in the continuous (−0.8±8.8 ms, t(18) = −0.24, p_fdr_ = .714, d_z_ = −0.056, BF_+0_ = 0.198) or apparent (1.5±6.7 ms, t(18) = 0.38, p_fdr_ = .545, d_z_ = 0.087, BF_+0_ = 0.326) conditions. In the static condition, responses were faster on different-than on same-object trials (−15.8±10.9 ms, t(18) = −2.55, d_z_ = −0.585). Our planned comparison was one-tailed in the predicted direction and cannot evaluate a difference of this sign (p_fdr_ = .990, BF_+0_ = 0.059). Treated as exploratory and tested two-tailed, the difference is reliable (p = .020, BF_10_ = 2.93, uncorrected).

Error rates did not follow the OSPB pattern. At *D_lon_*_g_, errors were 6.8% (SD = 3.8%) for same-, 7.1% (SD = 3.9%) for different-, and 6.7% (SD = 3.5%) for no-match trials; at *D_vsac_*, 6.3% (SD = 3.8%), 6.0% (SD = 2.9%), and 5.6% (SD = 3.7%), respectively.

### General Preview Benefit (RT_no-match_ – RT_different_)

At *D_lon_*_g_, significant GPBs were found in the static (11.4±9.2 ms, t(18) = 3.07, p_fdr_ = .013, d_z_ = 0.705, BF_10_ = 7.42) and continuous (10.3±8.8 ms, t(18) = 3.11, p_fdr_ = .013, d_z_ = 0.714, BF_10_ = 7.98) conditions, while the apparent condition did not reach significance (7.4±10.7 ms, t(18) = 1.64, p_fdr_ = .119, d_z_ = 0.376, BF_10_ = 0.736). At *D_vsac_*, significant GPBs were observed across all three motion conditions: static (11.5±9.3 ms, t(18) = 2.87, p_fdr_ = .015, d_z_ = 0.659, BF_10_ = 5.16), continuous (8.9±7.3 ms, t(18) = 3.20, p_fdr_ = .013, d_z_ = 0.735, BF_10_ = 9.43), and apparent (9.5±9.4 ms, t(18) = 2.40, p_fdr_ = .033, d_z_ = 0.550, BF_10_ = 2.27). These findings indicate that general preview benefits, independent of object identity, persist across both linking display durations (Figure 4B).

Since continuous motion never interrupts spatiotemporal continuity between preview and target, we also tested whether the continuous condition produced faster RTs than the apparent condition at each linking display duration using two-tailed paired t-tests. No significant differences (RT_continuous_ – RT_apparent_) were found at either *D_lon_*_g_ (6.5 ms, t(18) = 0.86, p_fdr_ = .403, d_z_ = 0.196) or *D_vsac_* (2.9 ms, t(18) = 0.89, p_fdr_ = .403, d_z_ = 0.204), in contrast to the significant slowing observed for the continuous condition at *D_lon_*_g_ in Experiment 1.

## Discussion

We found a robust OSPB at *D_lon_*_g_ for the continuous motion condition, replicating the key finding of Experiment 1. However, unlike Experiment 1, the static condition at long duration did not show a significant OSPB. We did not compare the two experiments directly, so this difference is equally consistent with sampling variability between independent groups; we return to it in the General Discussion.

Consistent with Experiment 1, no significant OSPBs were observed across any motion condition at *D_vsac_*, indicating that object-specific information is not conveyed at saccadic speeds, even when the objects carried a distinguishing feature. One limit on this conclusion concerns when the rings were visible. They appeared in the preview and target displays but not during the linking display, where they would have produced visible motion streaks, so the objects crossed that interval as bare, identical Gabor patches. The manipulation therefore asks whether a feature available at both endpoints supports object-specific transfer; it leaves open whether a feature that travels with the object would.

In the static condition, responses were faster on different-than on same-object trials, as in Experiment 1, though there the reversal emerged only at the shorter *D_vpeak_*. We return to this reversal in the General Discussion.

## General Discussion

Both experiments demonstrated robust OSPBs of around 20 ms at *D_lon_*_g_ for the continuous motion condition. In Experiment 1, the objects were identical Gabor patches, distinguishable only by their spatiotemporal trajectory, and object-specific information was nonetheless transferred across the motion. This extends the findings of Sasi et al. (2023), whose objects differed in shape and color, to objects defined by trajectory alone. Experiment 2 reproduced the benefit when a distinguishing surface feature present at preview and target.

In the static and apparent conditions, the previewed objects disappeared during the blank linking display and reappeared at the target display. For the static condition, where the objects stayed in the same place, this parallels the conditions that produce object substitution masking (Di Lollo et al., 2000; Enns & Di Lollo, 1997): provided the stimulus reappears in close spatial and temporal proximity, it can act as a trailing mask for the previewed representation, disrupting retrieval of object-specific information and potentially reversing the expected OSPB. Both requirements were met at our saccadic linking display durations in the static condition, but not in the apparent or continuous conditions, where the objects ended up 6 dva away and nothing followed at the previewed location. In agreement with this account, the OSPB reversed in the static condition at *D_vpeak_* in Experiment 1 and at *D_vsac_* in Experiment 2, and in both experiments the static benefit declined monotonically as the blank interval shortened.

The static condition at *D_lon_*_g_ is a separate matter. It produced a significant OSPB but not in Experiment 2, a difference we did not test directly, but which is consistent with sampling variability between independent groups. In any case, as noted above, the static OSPB is also consistent with a preview benefit tied to location rather than to the object, so its presence or absence bears less on object-specific transfer than the continuous condition does.

A further, complementary account specific to the reversal in the static condition concerns the detectability of the target display onset, which signals a participant that a response shall be given. In the static condition, the objects do not change location between preview and target, and at saccadic linking display durations the intervening blank lasts less than 40 ms. Under these conditions, the perceptual change signaling the onset of the target display is minimal. This is particularly the case in same-object trials, where the digit reappears on the same Gabor patch where it was already shown during the preview. In the different-object condition, by contrast, a digit appears on an object where it has not been before, creating a more visible cue to target display onset. This difference between same– and different-object trials could cause participants to initiate their response slightly faster in the different-than in the same-object condition. This onset-detection confound would not affect the continuous or apparent conditions, where the motion or apparent displacement itself provides an easily noticeable change. Future experiments could disambiguate these accounts by consistently signaling target display onset across conditions, for instance by adding a brief luminance change in the fixation point or an auditory signal at target display onset.

Critically, we did not find OSPBs at saccadic durations in either experiment. Instead, both experiments showed robust GPBs there. This pattern is informative mainly for what it rules out. Participants still responded faster to a previewed digit, whichever object now contained it, so the previewed content is neither erased nor masked by the motion itself. What is lost is the assignment of the content to a particular object. This dissociation has a precedent in the trans-saccadic literature. Henderson & Anes (1994) separated object-file review, a benefit tied to a particular object, from type priming, a benefit that requires only that an identity has been seen. These correspond to our OSPB and GPB, respectively, and they showed that the two can come apart across eye fixations. They obtained object-specific benefits across actual saccades; we obtained none with visual motion that merely matched the speed of one. In the terms of Kahneman et al. (1992), we would say that object files survive at saccadic speeds; what fails is the link between a file and the object in the world it belongs to. The present data do not exclude the stronger possibility that no file is formed under these conditions and that only a type-level representations persist.

Why should a real saccade support object-specific transfer where motion at the same speed does not? One strong candidate is predictive remapping driven by efference copy signals: before and during a saccade, the motor command to move the eyes generates an internal copy of the movement signal that can, in principle, update the spatial coordinates of object representations to anticipate the retinal consequences of the eye movement (Golomb & Mazer, 2021). If object-specific updating across saccades depends on such a signal, its absence would explain why matching the speed of a saccade is not sufficient: our participants held fixation throughout, so the displacement was never accompanied by an efference copy, and correspondence had to be established from the retinal input alone. This is consistent with studies on trans-saccadic integration that find integration of orientation information across a saccade but not when the same retinal displacement is replayed during passive viewing (Ganmor et al., 2015). Remapping cannot be the general mechanism, however. Our objects at *D_lon_*_g_ moved while the eyes were fixating, with no efference copy available, yet object-specific benefits were robust. Whatever links features to objects during ordinary object motion must therefore have a speed limit of its own—one that a real saccade may circumvent through the extraretinal signal it provides.

The speed dependence we observed also aligns with the multiple object tracking literature. Holcombe (2009) identified a slow temporal limit on attentive tracking that lies far the limits of low-level motion perception. Holcombe & Chen (2012) showed that this limit reflects a resource-constrained attentional process, as tracking two objects produced a substantially lower speed limit than tracking one object. In our continuous condition, two objects moved simultaneously, so on this account the ceiling should be correspondingly low. The mechanism that binds features to objects in the object-reviewing paradigm may be subject to the same constraint, though our design localizes that boundary only to somewhere between 20 and 154 deg/s.

Our findings in Experiment 2, showing that distinguishing features (rings outlining the Gabors) do not lead to OSPBs at saccadic durations, could contribute to a longstanding debate about the role of spatiotemporal continuity and surface features in object correspondence (Mitroff & Alvarez, 2007; Moore et al., 2010; Odic et al., 2012). Mitroff and Alvarez (2007) showed that spatiotemporal information, rather than surface features, is the main contributing factor in establishing object files. In contrast, more recent work has shown that surface features can influence object correspondence when spatiotemporal information is ambiguous (Moore et al., 2010, 2020). It may matter which surface feature is involved, not only whether one is present: Gordon & Vollmer (2010) found that colour enters an object’s episodic representation whether or not it is diagnostic of identity, but that changing it disrupts object continuity only for objects with a diagnostic colour; for letters and for objects without a characteristic colour, the object-specific benefit survived a colour change intact. The rings in Experiment 2 are non-diagnostic in exactly this sense, distinguishing two tokens without contributing to the identity of either, so on their account the rings would not be expected to support object-specific transfer. Our findings also align with Mitroff and Alvarez (2007): adding surface-feature information was insufficient to enable OSPBs. One further qualifications applies, however: The feature was not visible during the motion. Whether a feature that travelled with the object, or one that identified it, would behave differently remains open.

We did not expect OSPBs in the apparent condition, because the correspondence between preview and target objects is unspecified there. Odic, Roth and Flombaum (2012) showed that apparent motion can support object-file updating and produce significant OSPBs, but their benefits appeared only in a motion quartet whose geometry made one correspondence shorter than the other; with an equidistant quartet, the benefits were lost. Our arrangement was also equidistant, so each previewed object was equally likely to be assigned to either target location, presumably at random across trials. Accordingly, we found no OSPB in the apparent condition at any linking display duration in either experiment. The condition thus served its intended role as a control in which displacement occurs without a specified correspondence, rather than as a test of apparent motion as such.

Taken together, our findings suggest that the visual system retains preview information at saccadic speeds, as the consistent general preview benefits show, but cannot reliably link that information to specific objects. This link appears to require continuous motion slow enough to be tracked. Adding a distinguishing surface feature in Experiment 2 did not restore it. How that link is established and maintained, and what predictive oculomotor signals contribute to it, remains open.

## Acknowledgements

This research was supported by the European Research Council under the European Union’s Horizon 2020 research and innovation program (Grant 865715), and the Deutsche Forschungsgemeinschaft (grants RO3579/8-1, RO3579/10-1). The authors declare no competing interests. We thank Leonie Nitsche, Siri Tappesser, and Antonia Keller for their support with the data collection.

## Notes

https://osf.io/3zj6v/

https://osf.io/wjp5s/

### Competing Interest Statement

The authors have declared no competing interest.

## References

1. Balsdon, T., Schweitzer, R., Watson, T. L., & Rolfs, M. (2018). All is not lost: Post-saccadic contributions to the perceptual omission of intra-saccadic streaks. Consciousness and Cognition, 64, 19–31. 10.1016/j.concog.2018.05.004

2. Benjamini, Y., & Hochberg, Y. (1995). Controlling the False Discovery Rate: A Practical and Powerful Approach to Multiple Testing. Journal of the Royal Statistical Society Series B: Statistical Methodology, 57(1), 289–300. 10.1111/j.2517-6161.1995.tb02031.x

3. Campbell, F. W., & Wurtz, R. H. (1978). Saccadic omission: Why we do not see a grey-out during a saccadic eye movement. Vision Research, 18(10), 1297–1303. 10.1016/0042-6989(78)90219-5

4. Castet, E. (2009). Perception of Intra-saccadic Motion. In U. J. Ilg & G. S. Masson (Eds.), Dynamics of Visual Motion Processing (pp. 213–238). Springer US. 10.1007/978-1-4419-0781-3_10

5. Castet, E., Jeanjean, S., & Masson, G. S. (2002). Motion perception of saccade-induced retinal translation. Proceedings of the National Academy of Sciences, 99(23), 15159–15163. 10.1073/pnas.232377199

6. Castet, E., & Masson, G. S. (2000). Motion perception during saccadic eye movements. Nature Neuroscience, 3(2), 177–183. 10.1038/72124

7. Collewijn, H., Erkelens, C. J., & Steinman, R. M. (1988). Binocular co-ordination of human horizontal saccadic eye movements. The Journal of Physiology, 404(1), 157–182. 10.1113/jphysiol.1988.sp017284

8. Di Lollo, V., Enns, J. T., & Rensink, R. A. (2000). Competition for consciousness among visual events: The psychophysics of reentrant visual processes. Journal of Experimental Psychology: General, 129(4), 481–507. 10.1037/0096-3445.129.4.481

9. Enns, J. T., & Di Lollo, V. (1997). Object Substitution: A New Form of Masking in Unattended Visual Locations. Psychological Science, 8(2), 135–139. 10.1111/j.1467-9280.1997.tb00696.x

10. Ganmor, E., Landy, M. S., & Simoncelli, E. P. (2015). Near-optimal integration of orientation information across saccades. Journal of Vision, 15(16), 8. 10.1167/15.16.8

11. Garćia-Pérez, M. A., & Peli, E. (2001). Intrasaccadic Perception. The Journal of Neuroscience, 21(18), 7313–7322. 10.1523/JNEUROSCI.21-18-07313.2001

12. Golomb, J. D., & Mazer, J. A. (2021). Visual Remapping. Annual Review of Vision Science, 7(1), 257–277. 10.1146/annurev-vision-032321-100012

13. Gordon, R. D., & Vollmer, S. D. (2010). Episodic representation of diagnostic and nondiagnostic object colour. Visual Cognition, 18(5), 728–750. 10.1080/13506280903004190

14. Henderson, J. M., & Anes, M. D. (1994). Roles of object-file review and type priming in visual identification within and across eye fixations. Journal of Experimental Psychology: Human Perception and Performance, 20(4), 826–839. 10.1037/0096-1523.20.4.826

15. Higgins, E., & Rayner, K. (2015). Transsaccadic processing: Stability, integration, and the potential role of remapping. *Attention, Perception*, & Psychophysics, 77(1), 3–27. 10.3758/s13414-014-0751-y

16. Holcombe, A. O. (2009). Seeing slow and seeing fast: Two limits on perception. Trends in Cognitive Sciences, 13(5), 216–221. 10.1016/j.tics.2009.02.005

17. Holcombe, A. O., & Chen, W.-Y. (2012). Exhausting attentional tracking resources with a single fast-moving object. Cognition, 123(2), 218–228. 10.1016/j.cognition.2011.10.003

18. Ince, M., Hübner, C., & Rolfs, M. (2025). Object continuity through invisible retinal motion at saccadic speed. Proceedings of the National Academy of Sciences, 122(47), e2518968122. 10.1073/pnas.2518968122

19. Kahneman, D., Treisman, A., & Gibbs, B. J. (1992). The reviewing of object files: Object-specific integration of information. Cognitive Psychology, 24(2), 175–219. 10.1016/0010-0285(92)90007-O

20. Kleiner, M., Brainard, D. H., & Pelli, D. G. (2007). What’s new in psychtoolbox-3. Perception, 36(14), 1–16.

21. Krekelberg, B. (2024). *Matlab Toolbox for Bayes Factor Analysis* (Version v3.0) [Computer software]. Zenodo. 10.5281/ZENODO.13744717

22. Mattler, U., & Fendrich, R. (2007). Priming by motion too rapid to be consciously seen. Perception & Psychophysics, 69(8), 1389–1398. 10.3758/BF03192954

23. Mattler, U., & Fendrich, R. (2010). Consciousness mediated by neural transition states: How invisibly rapid motions can become visible. Consciousness and Cognition, 19(1), 172–185. 10.1016/j.concog.2009.12.015

24. Miles, W. R. (1930). Ocular Dominance in Human Adults. The Journal of General Psychology, 3(3), 412–430. 10.1080/00221309.1930.9918218

25. Mitroff, S. R., & Alvarez, G. A. (2007). Space and time, not surface features, guide object persistence. Psychonomic Bulletin & Review, 14(6), 1199–1204. 10.3758/BF03193113

26. Moore, C. M., Stephens, T., & Hein, E. (2010). Features, as well as space and time, guide object persistence. Psychonomic Bulletin & Review, 17(5), 731–736. 10.3758/PBR.17.5.731

27. Moore, C. M., Stephens, T., & Hein, E. (2020). Object correspondence: Using perceived causality to infer how the visual system knows what went where. *Attention, Perception*, & Psychophysics, 82(1), 181–192. 10.3758/s13414-019-01763-y

28. Nicolas, G., Kristensen, E., Dojat, M., & Guérin-Dugué, A. (2026). Eye movement dynamics are a key factor for intra-saccadic motion perception. Scientific Reports, 16(1), 8144. 10.1038/s41598-026-39420-8

29. Noles, N. S., Scholl, B. J., & Mitroff, S. R. (2005). The persistence of object file representations. Perception & Psychophysics, 67(2), 324–334. 10.3758/BF03206495

30. Odic, D., Roth, O., & Flombaum, J. I. (2012). The relationship between apparent motion and object files. Visual Cognition, 20(9), 1052–1081. 10.1080/13506285.2012.721405

31. Rolfs, M., & Schweitzer, R. (2022). Coupling perception to action through incidental sensory consequences of motor behaviour. Nature Reviews Psychology, 1(2), 112–123. 10.1038/s44159-021-00015-x

32. Rolfs, M., Schweitzer, R., Castet, E., Watson, T. L., & Ohl, S. (2025). Lawful kinematics link eye movements to the limits of high-speed perception. Nature Communications, 16(1), 3962. 10.1038/s41467-025-58659-9

33. Sasi, M., Friedman, S., & Lamy, D. (2023). The go no-go reviewing paradigm: A reliable method for measuring perceptual object-file updating. *Attention, Perception*, & Psychophysics, 85(1), 140–151. 10.3758/s13414-022-02606-z

34. Schweitzer, R., Doering, M., Seel, T., Raisch, J., & Rolfs, M. (2025). Saccadic omission revisited: What saccade-induced smear looks like. Psychological Review. 10.1037/rev0000574

35. Schweitzer, R., & Rolfs, M. (2020). Intra-saccadic motion streaks as cues to linking object locations across saccades. Journal of Vision, 20(4), 17. 10.1167/jov.20.4.17

36. Schweitzer, R., & Rolfs, M. (2021). Intrasaccadic motion streaks jump-start gaze correction. Science Advances, 7(30), eabf2218. 10.1126/sciadv.abf2218

37. Schweitzer, R., Seel, T., Raisch, J., & Rolfs, M. (2025). Early visual signatures and benefits of intra-saccadic motion streaks. PLOS Computational Biology, 21(9), e1013544. 10.1371/journal.pcbi.1013544

38. Stein, M., Fendrich, R., & Mattler, U. (2019). Stimulus dependencies of an illusory motion: Investigations of the Motion Bridging Effect. Journal of Vision, 19(5), 13. 10.1167/19.5.13

39. Stein, M., Fendrich, R., & Mattler, U. (2020). Encoding Information From Rotations Too Rapid To Be Consciously Perceived as Rotating: A Replication of the Motion Bridging Effect on a Liquid Crystal Display. I-Perception, 11(3), 2041669520925111. 10.1177/2041669520925111

40. Van Opstal, A. J., & Van Gisbergen, J. A. M. (1987). Skewness of saccadic velocity profiles: A unifying parameter for normal and slow saccades. Vision Research, 27(5), 731–745. 10.1016/0042-6989(87)90071-X

